# cGMP signaling regulates context-dependent sensory valence in *C. elegans*

**DOI:** 10.64898/2026.07.28.741361

**Authors:** Ricardo F. Frausto, Navonil Banerjee, Michelle L. Castelletto, Ava E. Bignell, Breanna Walsh, Elissa A. Hallem

**Affiliations:** Department of Microbiology, Immunology, and Molecular Genetics, University of California, Los Angeles, Los Angeles, California, USA; Molecular, Cellular, and Integrative Physiology Interdepartmental PhD Program, University of California, Los Angeles, Los Angeles, California, USA; Molecular Biology Interdepartmental PhD Program, University of California, Los Angeles, Los Angeles, California, USA; UCLA-Caltech Medical Scientist Training Program, University of California, Los Angeles, Los Angeles, California, USA; Molecular Biology Institute, University of California, Los Angeles, Los Angeles, California, USA

**Author notes:** Department of Neuroscience, Pomona College, Claremont, California, USA. Corresponding author (EAH).

## Abstract

An animal’s response to chemosensory cues depends on the animal’s prior experience, internal state, or life stage. However, the molecular mechanisms that regulate sensory valence (*i.e.,* whether a chemosensory cue is attractive or repulsive) remain poorly understood. We investigated the mechanisms that specify sensory valence using the responses of the free-living nematode *Caenorhabditis elegans* to carbon dioxide (CO_2_). *C. elegans* exhibits highly flexible responses to CO_2_: well-fed animals are repelled by CO_2_, while both starved animals and well-fed animals raised under high CO_2_ conditions are attracted to CO_2_. Here, we show that CO_2_ attraction in animals raised at high CO_2_ requires a cGMP signaling pathway that involves the cGMP-dependent protein kinase EGL-4. This pathway does not regulate CO_2_ response in starved animals, indicating that the role of EGL-4 in mediating CO_2_ attraction depends on satiety state. Cultivation under high CO_2_ conditions leads to increased cGMP levels in the CO_2_-detecting BAG neurons, consistent with a specific requirement for EGL-4 in high-CO_2_-cultivated animals. We also show that EGL-4 regulates CO_2_ valence by altering neuropeptide expression in BAG. Our results indicate that sensory valence is established in a context-dependent manner at the level of the primary sensory neuron.

## Introduction

A central feature of animal survival is the ability to appropriately respond to sensory stimuli present in the environment. Across animal phyla, sensory responses are used to avoid predators or pathogens, find food and mates, and navigate toward favorable environmental conditions [1–4]. Many of these responses are flexible and depend on the animal’s internal state, prior experience, and environmental context [5–11]. For example, in humans, food odors are perceived as pleasant during times of hunger, but satiety decreases the pleasantness of these same food odors [12]. However, the molecular mechanisms that drive changes in sensory valence remain poorly understood.

A sensory cue that elicits flexible responses from many animals, including nematodes, is carbon dioxide (CO_2_). CO_2_ can signal the presence of food, pathogens, predators, hosts, or conspecifics [5, 13–15]. Moreover, nematodes and other small invertebrates experience large fluctuations in environmental CO_2_ levels; while atmospheric levels of CO_2_ are ∼0.04%, CO_2_ levels in soil or compost can exceed 10% [16–18]. Across many species, behavioral responses to CO_2_ are subject to context-dependent modulation such that CO_2_ can be attractive, repulsive, or neutral; this modulation likely reflects the need for animals to exhibit diverse responses to CO_2_ as they experience varied environments [5, 19–22].

The response of *C. elegans* to CO_2_ offers a powerful system for addressing the mechanisms that regulate sensory valence. In *C. elegans,* CO_2_ is repulsive for well-fed animals that were raised under normal atmospheric conditions (*i.e.,* low CO_2_) [23–25] but attractive for starved animals [26], animals raised under high CO_2_ conditions [27], dauer larvae [28, 29], and non-dauer developmentally arrested larvae [25]. CO_2_ response is mediated primarily by the CO_2_-detecting BAG neurons in the head, which are activated by upshifts in CO_2_ concentration [30]. Notably, increases in the CO_2_ level reliably lead to activation in BAG neurons, but can result in repulsive, neutral, and attractive downstream behavioral responses depending on an animal’s context [26, 27, 30, 31]. This juxtaposition suggests that previously unidentified molecular mechanisms modulate the transduction of directionally stable neural responses in the BAG sensory neurons into distinct, flexible behaviors.

To gain insight into the molecular basis of CO_2_ valence encoding, we investigated the signaling pathways that regulate CO_2_ response in *C. elegans* adults. We show that CO_2_ attraction in adults raised under high CO_2_ conditions requires a cGMP signaling pathway mediated by the cGMP-dependent protein kinase EGL-4 [32]. Although CO_2_ is similarly attractive to starved adults, EGL-4 is not required for CO_2_ attraction in starved adults. EGL-4 is also not required for CO_2_ repulsion in well-fed adults raised under low CO_2_ conditions. Thus, EGL-4 regulates CO_2_ valence in the context of prior CO_2_ exposure, but not in the context of hunger vs. satiety. EGL-4 acts within the BAG sensory neurons to modulate CO_2_ valence by regulating neuropeptide gene expression. Our results demonstrate that cGMP signaling in primary sensory neurons establishes CO_2_ valence in a context-dependent manner to drive flexible behavioral responses to CO_2_.

## Results

### EGL-4 regulates CO_2_ valence in animals cultured at high CO_2_ but not starved animals

The *C. elegans* BAG neurons detect CO_2_ using the receptor guanylate cyclase GCY-9 – a putative receptor for molecular CO_2_ – and the cGMP-gated calcium channel TAX-2/TAX-4 [23, 30, 33]. Since CO_2_ detection is mediated by a cGMP signaling pathway, we asked if proteins known to be regulated by cGMP signaling play a role in modulating CO_2_ response. One such protein is the cGMP-dependent protein kinase EGL-4, which is broadly expressed in *C. elegans* sensory neurons and is a key intracellular regulator of chemosensory behavior [32, 34–39]. Since *egl-4* is expressed in the BAG neurons of adult hermaphrodites [40], we investigated the possibility that it regulates CO_2_ response. We examined the CO_2_-evoked behavior of young adults using a CO_2_ chemotaxis assay in which animals were allowed to navigate freely in a CO_2_ gradient, enabling us to quantify their attractive, neutral, or aversive responses to CO_2_ (Fig S1). We first confirmed that well-fed animals raised at low CO_2_ (*i.e.,* ambient CO_2_) were repelled by CO_2_, whereas starved animals raised at low CO_2_ and well-fed animals raised at high CO_2_ (*i.e.,* 2.5% CO_2_) were attracted to CO_2_ (Fig 1A). To examine the requirement for *egl-4* in mediating CO_2_ response, we then compared the behavior of wild-type vs. *egl-4* loss-of-function (*lof*) animals under the same conditions. We found that both wild-type and *egl-4(lof)* well-fed animals raised at low CO_2_ were repelled by CO_2_, and both wild-type and *egl-4(lof)* starved animals were attracted to CO_2_ (Fig 1B and S2). However, whereas well-fed wild-type animals raised at high CO_2_ were attracted to CO_2_, well-fed *egl-4(lof)* animals raised at high CO_2_ were repelled by CO_2_ (Fig 1B and S2). These results suggest that EGL-4 is not required for the valence switch from CO_2_ aversion to attraction that occurs during starvation; in contrast, EGL-4 is required for the CO_2_ valence switch from aversion to attraction that occurs in animals raised under high CO_2_ conditions.

**Fig 1.**
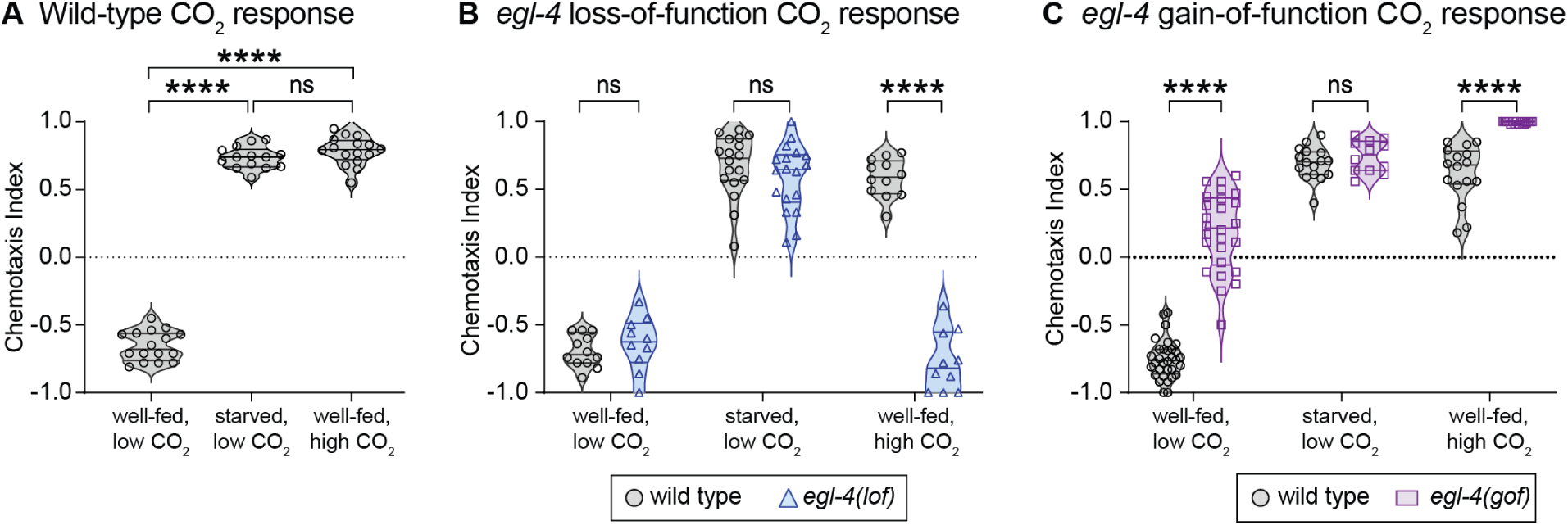
EGL-4 mediates attraction to CO_2_ in animals raised at high CO_2_ but not starved animals. (**A**) Chemotaxis of wild-type animals across conditions. \*\*\*\**p*<0.0001, ns = not significant, one-way ANOVA with Tukey’s post-test. n = 14-16 trials per condition. (**B**) Chemotaxis of wild-type animals vs. animals containing the loss-of-function (*lof*) allele *egl-4(n479)* across conditions. \*\*\*\**p*<0.0001, ns = not significant, two-way ANOVA with Šidák’s post-test. n = 10-18 trials per condition. **(C)** Chemotaxis of wild-type animals vs. animals containing the gain-of-function (*gof*) allele *egl-4(mg410)* across conditions. \*\*\*\**p*<0.0001, ns = not significant, two-way ANOVA with Šidák’s post-test. n = 16-32 trials per condition. For all graphs, symbols represent individual trials and lines indicate medians and interquartile ranges.

We also examined animals with a gain-of-function (*gof*) mutation in *egl-4* consisting of a point mutation that enables EGL-4 to autophosphorylate independent of cellular cGMP level, leading to constitutive activity [34]. We found that whereas well-fed wild-type animals raised at low CO_2_ are repelled by CO_2_, well-fed *egl-4(gof)* animals raised at low CO_2_ are weakly attracted or neutral to CO_2_ (Fig 1C). Moreover, well-fed *egl-4(gof)* animals raised at high CO_2_ exhibit more robust CO_2_ attraction than wild-type animals (Fig 1C). These results further suggest that increased EGL-4 activity in well-fed animals promotes a valence shift from CO_2_ aversion to CO_2_ attraction. In contrast, starved *egl-4(gof)* animals showed normal CO_2_ attraction, confirming that EGL-4 activity does not impact the valence switch from CO_2_ aversion to CO_2_ attraction that occurs in response to starvation (Fig 1C).

### EGL-4 acts in BAG sensory neurons to promote CO_2_ attraction

To determine where in the CO_2_ circuit EGL-4 acts to regulate CO_2_ response, we performed a rescue experiment in which a wild-type copy of the *egl-4* gene was restored to *egl-4(lof)* mutants in either all EGL-4-positive cells, all neurons, or the subset of sensory neurons that express the cGMP-gated cation channel subunit TAX-4 [41]. Rescue of *egl-4* in each case restored CO_2_ attraction in well-fed *egl-4(lof)* animals raised at high CO_2_ (Fig 2A), indicating that EGL-4 acts in TAX-4-expressing neurons to establish CO_2_ attraction. The CO_2_-detecting BAG neurons are one of several pairs of head sensory neurons that express TAX-4 [41–43]. To determine if EGL-4 acts in BAG or one of the other TAX-4-expressing neurons to regulate CO_2_ response, we restored wild-type *egl-4* expression in either single pairs or small subsets of TAX-4+ neurons (Fig 2B). We found that rescue solely in the BAG neurons was sufficient to restore CO_2_ attraction to well-fed *egl-4(lof)* animals raised at high CO_2_ (Fig 2B). Thus, EGL-4 acts in BAG neurons to establish CO_2_ attraction in animals raised at high CO_2_.

**Fig 2.**
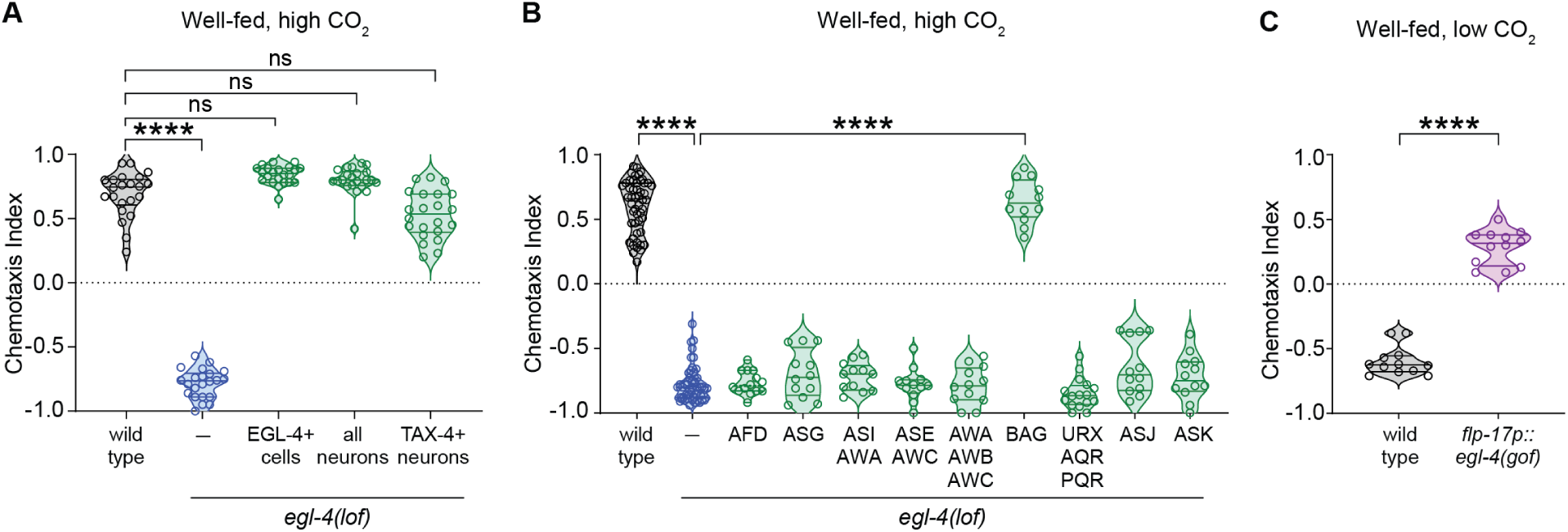
EGL-4 acts in BAG neurons to mediate the valence switch to CO_2_ attraction that occurs following cultivation at high CO_2_. **(A)** Chemotaxis of wild-type animals, *egl-4(lof)* animals, and *egl-4(lof)* animals in which wild-type *egl-4* expression was restored in either all EGL-4-positive cells, all neurons, or all TAX-4-positive neurons (*lof* allele = *ky185*). Animals were well-fed and raised at high CO_2_. \*\*\*\**p*<0.0001, ns = not significant, Kruskal-Wallis test with Dunn’s post-test. n = 22 trials per condition. **(B)** Chemotaxis of wild-type animals, *egl-4(lof)* animals, and *egl-4(lof)* animals in which wild-type *egl-4* expression was restored in different subsets of TAX-4+ sensory neurons (*lof* allele = *ky185*). \*\*\*\**p*<0.0001, Kruskal-Wallis test with Dunn’s post-test. n = 12-50 trials per genotype. **(C)** Chemotaxis of wild-type animals and animals expressing the *egl-4(mg410)* gain-of-function allele specifically in the BAG neurons. \*\*\*\**p*<0.0001, Welch’s t-test. N = 12 trials per genotype. For all graphs, symbols represent individual trials and lines indicate medians and interquartile ranges.

To determine whether EGL-4 activity in BAG is sufficient to drive a valence switch from CO_2_ aversion to CO_2_ attraction, we examined the CO_2_ response of well-fed animals cultivated under low CO_2_ conditions that expressed a BAG-specific *egl-4(gof)* allele. We found that despite being well-fed and raised at low CO_2_, transgenic animals expressing the BAG-specific *egl-4(gof)* allele were attracted to CO_2_ (Fig. 2C). Thus, increased EGL-4 activity in BAG is sufficient to switch CO_2_ response from repulsive to attractive in well-fed animals.

### Elevated cGMP levels in BAG promote CO_2_ attraction in animals raised at high CO_2_

To gain insight into why EGL-4 is required for CO_2_ attraction specifically in animals raised at high CO_2_, we investigated the possibility that tonic exposure to high CO_2_ leads to elevated cGMP levels in BAG, resulting in increased EGL-4 activity. It was previously shown that in addition to responding to acute increases in CO_2_ levels [30, 33, 44], the BAG neurons show tonic activity in response to prolonged CO_2_ exposure [44]. Moreover, expression of the putative CO_2_ receptor GCY-9 – a receptor guanylate cyclase that, when activated, mediates the conversion of GTP into cGMP [30, 33, 45] – is upregulated by prolonged CO_2_ exposure [27]. Thus, in animals raised under high CO_2_ conditions, tonic CO_2_ exposure could lead to elevation of cGMP levels in BAG via activation of GCY-9. This in turn could result in increased EGL-4 activity in BAG, since EGL-4 activity is cGMP-dependent [46]. To test this possibility, we imaged cGMP levels in the BAG neurons of well-fed animals raised at low CO_2_, starved animals raised at low CO_2_, and well-fed animals raised at high CO_2_ using the genetically encoded cGMP sensor FlincG3 [47]. We found that high-CO_2_ culturing, but not starvation, resulted in elevated cGMP levels in BAG (Fig 3A-B). These results suggest that CO_2_ attraction in animals cultured at high CO_2_ may be conferred by tonically elevated cGMP levels, which in turn increase and maintain tonic EGL-4 activity.

**Fig 3.**
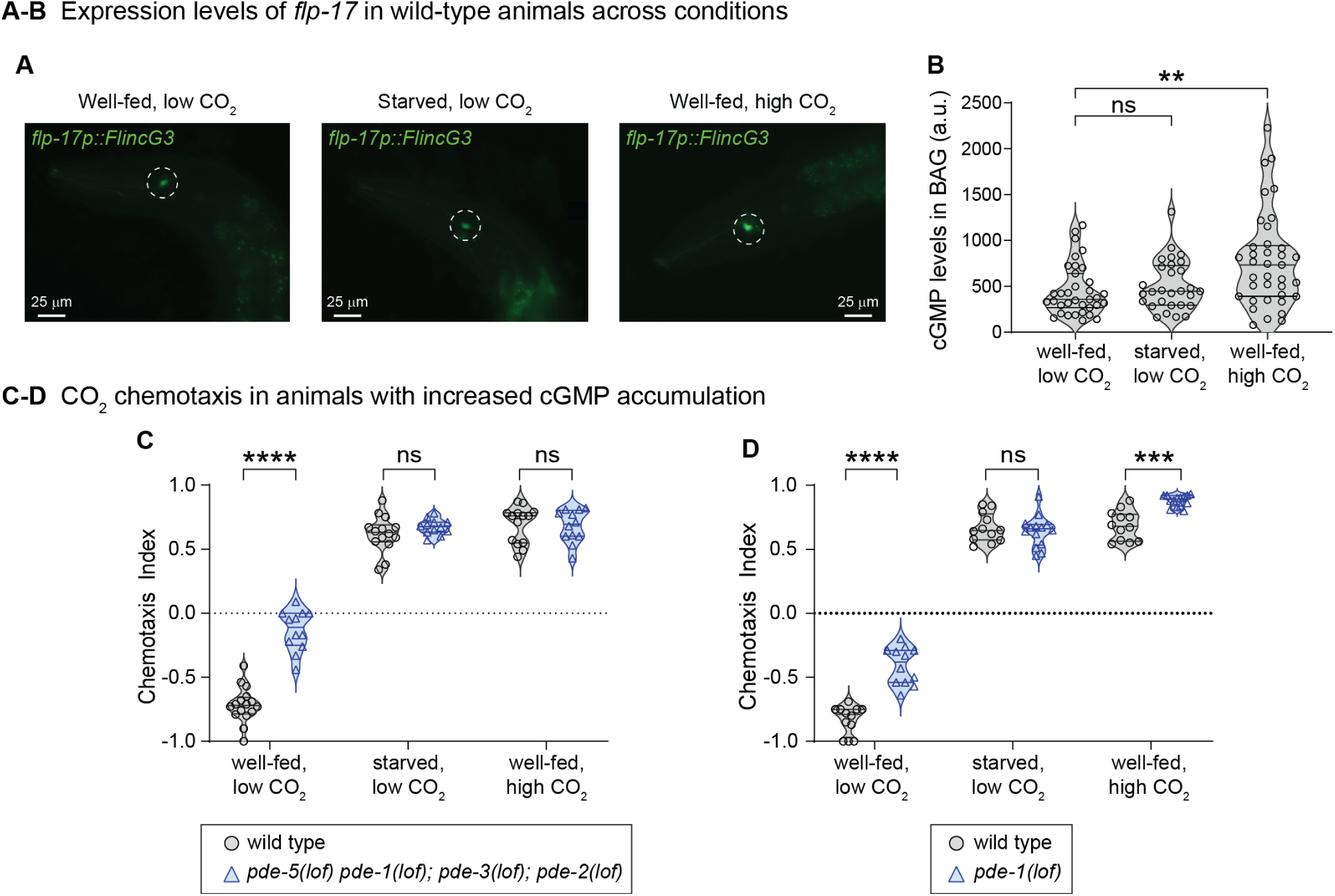
Elevated cGMP signaling promotes CO_2_ attraction. **(A)** Representative epifluorescence images of cGMP levels in BAG neurons across conditions. cGMP levels in individual BAG neuron cell bodies were visualized using the genetically encoded cGMP sensor FlincG3 [47]. **(B)** Quantification of cGMP levels in BAG neurons across conditions. \*\**p*<0.01, ns = not significant, Kruskal-Wallis test with Dunn’s post-test. n = 29-37 animals for each condition. In the graph, each symbol represents the fluorescence intensity of a single BAG neuron; lines indicate medians and interquartile ranges. **(C)** Chemotaxis of wild-type animals vs. *pde-5(nj49) pde-1(nj57); pde-3(nj59); pde-2(nj58)* quadruple mutants, which lack the four phosphodiesterases predicted to hydrolyze cGMP. \*\*\*\**p*<0.0001, ns = not significant, two-way ANOVA with Šidák’s post-test. n = 12-16 trials per genotype and condition. **(D)** Chemotaxis of wild-type animals vs. *pde-1(nj57)* mutants. \*\*\**p*<0.001, \*\*\*\**p*<0.0001, ns = not significant, two-way ANOVA with Šidák’s post-test. N = 12-14 trials per genotype and condition. For C-D, symbols represent individual trials and lines indicate medians and interquartile ranges.

To further investigate whether elevated cGMP levels promote CO_2_ attraction, we examined the CO_2_ response of animals that lack phosphodiesterases, resulting in the intracellular accumulation of cGMP. The *C. elegans* genome encodes six phosphodiesterases, four of which are predicted to hydrolyze cGMP: PDE-1, PDE-2, PDE-3, and PDE-5 [48]. We first tested a quadruple mutant that lacks all four of these phosphodiesterases [49]. We found that well-fed quadruple mutants raised at low CO_2_ show severely reduced CO_2_ repulsion relative to wild-type animals, consistent with elevated cGMP levels promoting a valence shift away from CO_2_ repulsion (Fig 3C). We then focused on *pde-1*, since it is expressed in the sensory endings of BAG neurons and regulates neuropeptide expression in BAG and CO_2_-evoked turning [45, 50]. We found that well-fed *pde-1* animals raised at low CO_2_ show reduced CO_2_ repulsion relative to wild-type animals, whereas well-fed *pde-1* animals raised at high CO_2_ show enhanced CO_2_ attraction relative to wild-type animals (Fig 3D). In contrast, starved wild-type and *pde-1(lof)* animals showed comparable responses to CO_2_ (Fig 3D). We note that unlike the *pde-1* mutants raised at high CO_2_, the quadruple mutants raised at high CO_2_ showed normal CO_2_ attraction, which could result from more widespread changes in cyclic nucleotide signaling in these mutants. Together, these results are consistent with a model in which elevated tonic cGMP levels in BAG promote CO_2_ attraction in animals raised at high CO_2_.

### EGL-4 is not required for the CO_2_-evoked calcium activity of BAG neurons

BAG neurons show robust CO_2_-evoked depolarizations in well-fed animals raised at low CO_2_, well-fed animals raised at high CO_2_, and starved animals raised at low CO_2_ [26, 27, 30]. To determine whether EGL-4 is required for CO_2_-evoked calcium responses in BAG, we performed calcium imaging on *egl-4(lof)* mutants. We found that the BAG neurons of *egl-4(lof)* mutants show normal depolarizing calcium responses to CO_2_ (Fig 4). Larger calcium responses were observed in animals raised at high CO_2_ (Fig 4), as previously described and consistent with the increased *gcy-9* expression in high-CO_2_-reared animals [27]. However, no significant differences were observed in the level of calcium influx between wild-type and *egl-4(lof)* animals (Fig 4). Thus, EGL-4 regulates CO_2_ valence but is not required for CO_2_-evoked calcium influx within BAG neurons.

**Fig 4.**
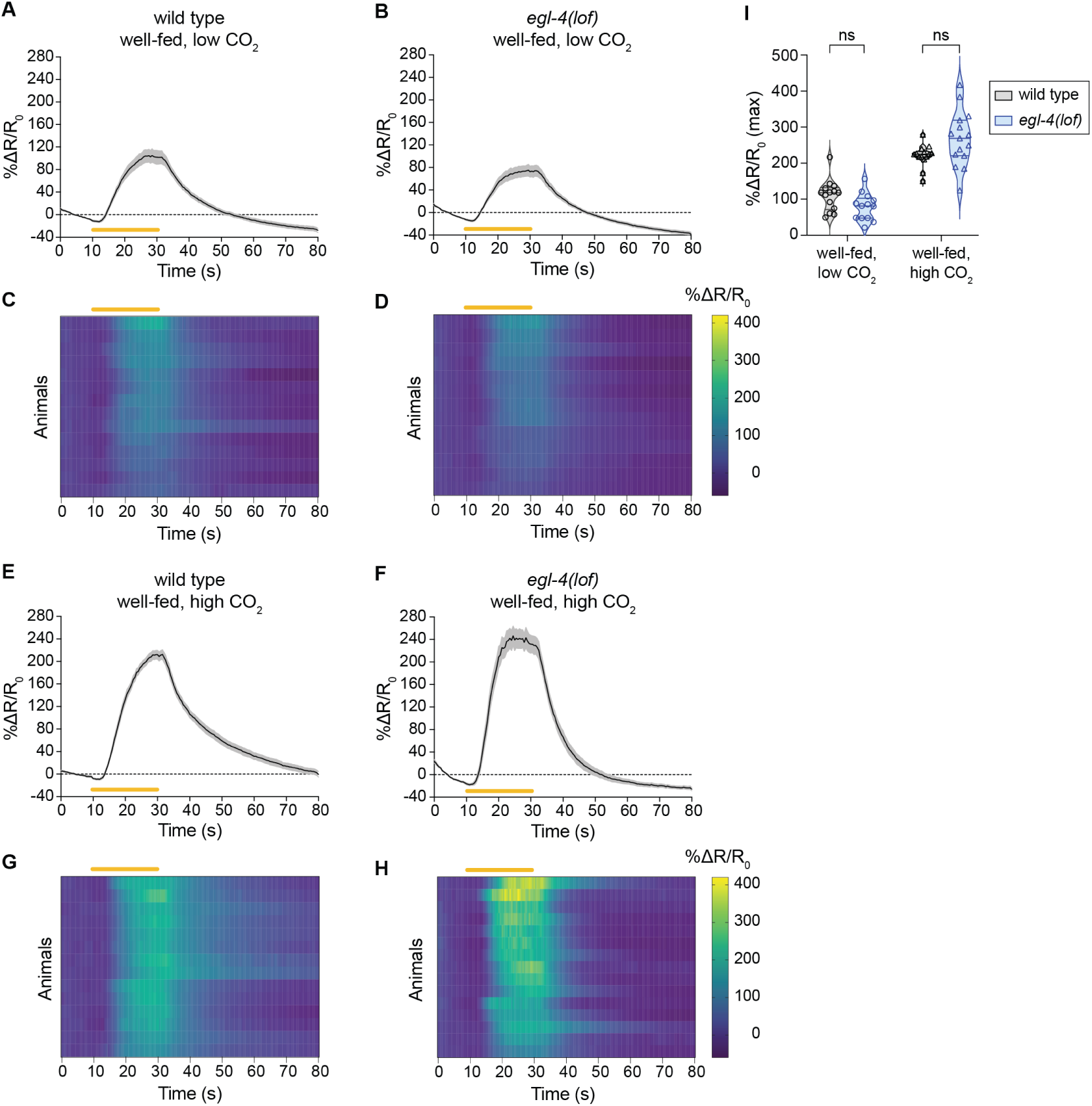
The BAG neurons of *egl-4(lof)* mutants respond normally to an acute CO_2_ stimulus. **(A-D)** Calcium responses of the BAG neurons of wild-type vs. *egl-4(lof)* mutants that were well-fed and raised at low CO_2_ to an acute CO_2_ stimulus. The genetically encoded ratiometric calcium indicator yellow cameleon YC3.60 was used to measure calcium responses in BAG neurons [73]. Graphs show calcium transients in the BAG neurons of wild-type (**A**) vs. *egl-4(lof)* (**B**) animals (*lof* allele = *ky185*). Lines show mean calcium responses and gray shading indicates SEM. Orange lines indicate the timing and duration of the CO_2_ pulse. Heatmaps show the BAG calcium responses of the individual wild-type (**C**) or *egl-4(lof)* (**D**) animals shown in A-B. Each row represents the response of an individual animal. Response magnitudes are color-coded according to the % ΔR/R_0_ gradient scale shown to the right. Orange lines indicate the timing and duration of the CO_2_ pulse, and rows are ordered by hierarchical cluster analysis. n = 13-14 animals per genotype. **(E-H)** Calcium responses of the BAG neurons of wild-type vs. *egl-4(lof)* mutants that were well-fed and raised at high CO_2_ to an acute CO_2_ stimulus. Graphs show calcium transients in the BAG neurons of wild-type (**E**) vs. *egl-4(lof)* (**F**) animals (*lof* allele = *ky185*). Heatmaps show the BAG calcium responses of the individual wild-type (**G**) or *egl-4(lof)* (**H**) animals shown in E-F. Parameters are as specified above. n = 14-15 animals per genotype. (**I**) Quantification of maximum calcium responses across conditions. Each data point represents the response of a single animal shown in A-H. n = 13-15 animals per genotype and condition. ns = not significant, two-way ANOVA with Tukey’s post-test; only adjacent comparisons across genotypes are shown. In the graph, symbols represent the maximum % ΔR/R_0_ for each neuron and lines indicate medians and interquartile ranges.

### EGL-4 regulates CO_2_-evoked behavior by modulating neuropeptide gene expression

We next investigated the downstream mechanisms by which EGL-4 promotes CO_2_ attraction in animals raised under high CO_2_ conditions. The FMRFamide-like neuropeptide FLP-17 is expressed in BAG and drives CO_2_-evoked behavior [27]. To test whether the effect of EGL-4 on CO_2_ attraction is mediated by FLP-17, we quantified *flp-17* expression in the BAG neurons of wild-type vs. *egl-4(lof)* animals using a strain that expresses GFP from the endogenous *flp-17* locus (G. Valperga and O. Hobert, in preparation). We found that *flp-17* expression is reduced in *egl-4(lof)* animals raised under both low and high CO_2_ conditions (Fig 5A-B). However, while we observed residual *flp-17* expression in *egl-4(lof)* animals raised under low CO_2_ conditions, *flp-17* expression was almost completely eliminated in high-CO_2_-cultivated *egl-4(lof)* animals (Fig 5A-B). These results suggest that EGL-4 promotes CO_2_ attraction in high-CO_2_-cultivated animals by regulating *flp-17* expression in BAG. To further test this possibility, we overexpressed *flp-17* in BAG in *egl-4(lof)* animals using the BAG-specific *gcy-33* promoter. We found that the loss of CO_2_ attraction in *egl-4(lof)* animals raised under high-CO_2_ conditions was partially rescued by *flp-17* overexpression in BAG (Fig 5C). Thus, EGL-4 promotes CO_2_ attraction in high-CO_2_-cultivated animals at least in part by regulating neuropeptide gene expression.

**Fig 5.**
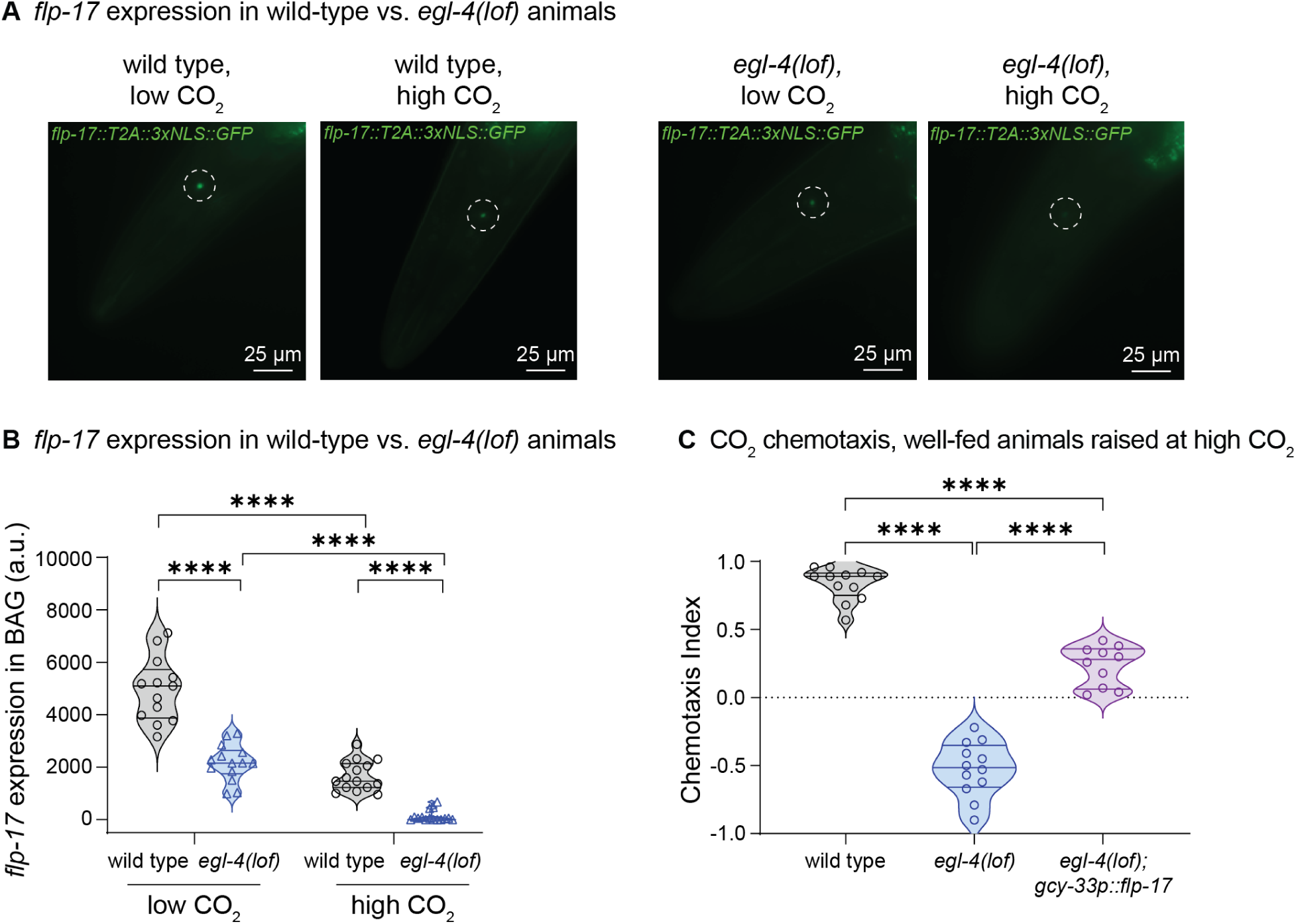
FLP-17 neuropeptides act downstream of EGL-4 to mediate CO_2_ attraction. (**A**) Representative epifluorescence images of nuclear GFP in BAG neurons of well-fed *flp-17(syb3184)* and *egl-4(lof); flp-17(syb3184)* animals raised at high CO_2_ (*egl-4*(*lof*) allele = *ky185*). The endogenously tagged *flp-17(syb3185)* animals produce 3xNLS-GFP and FLP-17 as separate proteins from a single transcript using a T2A self-cleaving peptide sequence (G. Valperga and O. Hobert, in preparation). (**B**) Quantification of nuclear 3xNLS-GFP levels in BAG as an indicator of *flp-17* expression. \*\*\*\**p*<0.0001, Kruskal-Wallis test with Dunn’s post-test. n = 13-15 neurons per genotype and condition. In the graph, each symbol represents a single BAG neuron; lines indicate medians and interquartile ranges. (**C**) Chemotaxis of well-fed wild-type, *egl-4(lof)*, and *egl-4(lof); gcy-33p::flp-17* animals raised at high CO_2_. The *egl-4(lof); gcy-33p::flp-17* animals overexpress *flp-17* specifically in BAG. \*\*\*\**p*<0.0001, one-way ANOVA with Tukey’s post-test. n = 10-12 trials per condition. In the graph, symbols represent individual trials and lines indicate medians and interquartile ranges.

## Discussion

Here, we show that cGMP signaling acts in the CO_2_-detecting BAG neurons to establish CO_2_ valence in *C. elegans*. CO_2_ response in *C. elegans* adults is highly context-dependent, such that CO_2_ is repulsive for well-fed animals raised at low CO_2_ but attractive for starved animals raised at low CO_2_ or well-fed animals raised at high CO_2_ [23, 24, 26, 27, 30, 44]. Moreover, CO_2_ valence is flexible and can switch from attractive to repulsive and vice versa [26, 27]. We show that in the CO_2_-detecting BAG neurons, increased cGMP signaling promotes the shift from CO_2_ repulsion to attraction in well-fed animals raised at high CO_2_ via the cGMP-dependent protein kinase EGL-4. In contrast, EGL-4 does not regulate CO_2_ response in starved animals. Thus, distinct mechanisms regulate CO_2_ valence in *C. elegans* adults depending on the context.

cGMP signaling has previously been shown to modulate *C. elegans* sensory behavior in other contexts. For example, cGMP signaling via EGL-4 modulates ASH-mediated nociceptive behavior in *C. elegans* by phosphorylating the regulators of G protein signaling RGS-2 and RGS-3 [37, 51]. It also causes the AWC-mediated switch from butanone attraction to repulsion that occurs when starved animals are exposed to a prolonged butanone stimulus [52]. Here, we demonstrate a new role for cGMP signaling in regulating CO_2_ valence in the BAG neurons. Our work supports a model in which prolonged exposure to elevated CO_2_ levels leads to tonic activation of GCY-9 in BAG [30], resulting in elevated levels of cGMP and activation of EGL-4, which promotes CO_2_ attraction (Fig 6).

**Fig 6.**
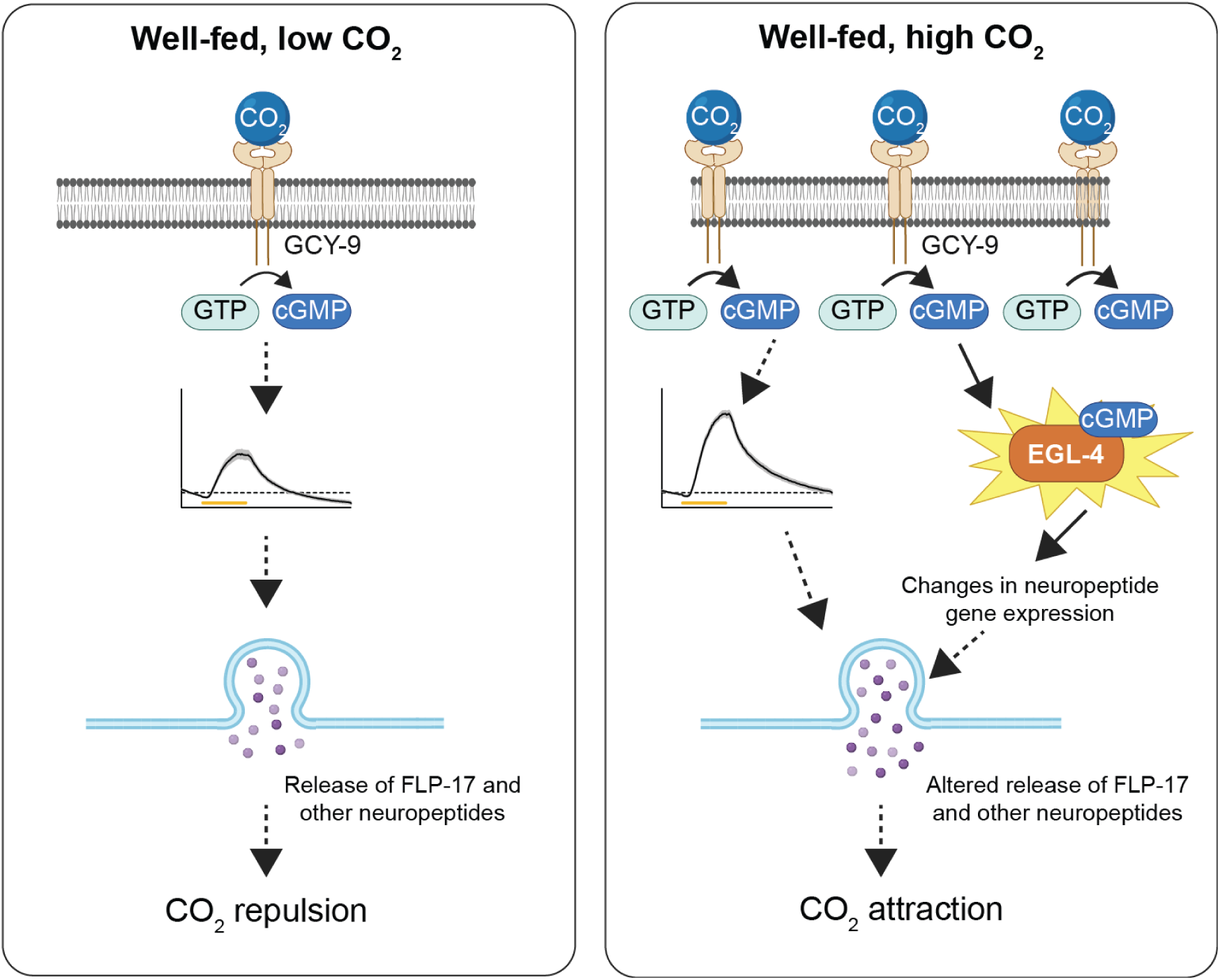
A model for CO_2_ valence determination in the CO_2_-detecting BAG neurons of well-fed animals exposed to low vs. high CO_2_ conditions. Under low CO_2_ conditions, an acute increase in CO_2_ response activates the receptor guanylate cycle GCY-9, resulting in cGMP production, BAG depolarization, and release of FLP-17 and other neuropeptides from BAG that drive a repulsive response to CO_2_ (left). Under high CO_2_ conditions, prolonged exposure to high CO_2_ results in increased expression of GCY-9 and increased GCY-9 signaling, resulting in increased cGMP production. This results in increased BAG depolarization and activation of the cGMP-dependent protein kinase EGL-4. EGL-4 promotes changes in expression of *flp-17* and likely other neuropeptide genes, resulting in altered neuropeptide release from BAG that drives an attractive response to CO_2_ (right).

We further show that EGL-4 promotes CO_2_ attraction by regulating *flp-17* expression (Fig 6). FLP-17 is one of several neuropeptides previously shown to play a role in CO_2_ response [27, 29]. A previous study found that expression of some neuropeptides in BAG is dependent on tonic BAG activity and the resulting elevation of cGMP levels [45]. Together with our results, this suggests a model in which CO_2_ valence is determined by context-dependent changes in the neuropeptide repertoire of BAG, resulting in differential activation of downstream interneurons [27]. The BAG neurons express over a dozen different neuropeptides that are predicted to act on over 50 neuron classes [53]; thus, context-dependent changes in neuropeptide expression in BAG could have dramatic effects on CO_2_ circuit function. While EGL-4 regulates expression of *flp-17* – and likely other neuropeptides – in animals cultivated at high CO_2_, the factors that regulate neuropeptide expression in starved animals and well-fed animals cultivated at low CO_2_ remain to be determined.

Our results pinpoint where valence is encoded in the CO_2_ circuit. We previously found that BAG shows a CO_2_-evoked increase in calcium levels regardless of the valence of the CO_2_ stimulus, whereas CO_2_-microcircuit interneurons show distinct activity patterns depending on the context [26, 27]. These results suggested that CO_2_ valence is encoded either in the BAG neurons downstream or independent of their calcium response or in the downstream interneurons. Our finding that EGL-4 acts in BAG to regulate CO_2_ response indicates that CO_2_ valence is encoded in the BAG neurons. However, it is possible that context-dependent changes in BAG signaling are complemented by changes in neuropeptide or neurotransmitter receptor expression in downstream interneurons.

CO_2_ valence encoding is reminiscent of valence encoding within the *C. elegans* thermosensory and gustatory circuits. In the thermosensory circuit, the AFD thermosensory neurons are activated by temperature increases regardless of response valence but opposing glutamatergic and neuropeptidergic signals from AFD evoke distinct activity patterns in downstream interneurons that are valence-dependent [54]. Similarly, in the gustatory circuit, the ASER gustatory neurons are activated by sodium chloride regardless of response valence but evoke valence-dependent activity patterns in downstream interneurons that are determined by basal glutamate level in ASER [11]. This circuit design, in which the same neurons mediate responses of opposite valence by altering their presynaptic output, contrasts with the more well-known mechanism of valence encoding in which distinct pathways mediate appetitive vs. aversive responses [55–60]. Similar circuit designs have been found in mammals, where the corticotropin-releasing factor (CRF)-releasing neurons of the paraventricular nucleus of the hypothalamus are differentially activated by appetitive vs. aversive stimuli to drive appropriate behavioral responses [61]. Thus, circuits in which the same neurons mediate responses of opposite valence through context-dependent modulation of their presynaptic output may be more common than previously recognized. Our results highlight the utility of *C. elegans* for decoding the functional architecture of these circuits.

## Methods

### C. elegans culturing

*C. elegans* young adult hermaphrodites were used for all experiments. Strains were maintained at 15°C and atmospheric CO_2_ (∼0.04% CO_2_) on 6-cm Petri plates containing 2% nematode growth media (NGM) with a lawn of *Escherichia coli* OP50. Animals used for experiments were maintained at room temperature (∼23°C) for at least three days prior to performing experiments. To ensure animals were maintained well-fed, especially those with strong roaming/dispersal phenotypes, some plates were seeded with 350 µL of concentrated OP50, which was spread to cover as much of the surface area of the plate as possible. Concentrated OP50 was prepared as previously described with minor modifications [62, 63]. Briefly, a single colony of OP50 from a streak plate was used to inoculate 5 mL of Luria Broth (LB) that was then incubated at 37°C in a shaker at 225 rpm for 4-6 h. This was used to inoculate 500 mL of LB, which was then incubated overnight (12-14 h) with shaking. The culture was centrifuged at 3000 *x g* for 15 min. Supernatant was discarded, and the pellet was resuspended in 100 mL of LB. This was stored at 4°C for no longer than 1 month.

### *C. elegans* crosses and genotyping

A list of strains used is provided in Table S1. For all experiments, *C. elegans* N2 Bristol was used as a control. EAH501 *egl-4(ky185); flp17(syb3184[flp-17::T2A::3xNLS::GFP])* was generated by crossing PHX3184 *flp-17(syb3184[flp-17::T2A::3xNLS::GFP])* with the *egl-4(ky185)* strain. EAH417 *egl-4(ky185); dbEx[flp-17p::YC3.60, lin-15(+)]* was made by crossing AX2073 *lin-15(n765ts); dbEx[flp-17p::YC3.60, lin-15(+)]* with the *egl-4(ky185)* strain. Genotyping was performed with Platinum Taq (ThermoFisher Scientific) or GoTaq (Promega) using the following reaction conditions: initial denaturation at 94°C for 2 min; cycled 35x through 94°C denaturation for 30 s, 55°C annealing for 30 s, and 72°C extension (time based on 1 kb/min); final extension at 72°C for 5 min. Primer sequences and amplicon sizes are listed in Table S2.

### Molecular biology and generation of transgenic animals

Transgenic animals carrying the transgene on an extrachromosomal array were generated using a standard intragonadal microinjection protocol [64]. DNA mixtures were prepared with a total plasmid DNA concentration of 100 ng/µL, of which 50 ng/µL (unless otherwise noted) consisted of the constructed plasmid and the remainder was a filler plasmid (pBluescript) or a plasmid containing *unc-122p::DsRed*, which labels coelomocytes and was used as a co-injection marker (Addgene plasmid #8938, a gift from P. Sengupta) [65]. Multiple lines of each transgenic strain were generated, and at least two independent lines of each genotype were tested.

Plasmids for neuron-specific rescue of *egl-4* (isoform 2a.1) were generated as follows. All rescue constructs were injected into *egl-4(ky185)* animals.

#### BAG

For cell-specific expression in BAG, the plasmid *flp-17p::egl-4.2a.1::SL2::GFP* (pRFF001) was generated from *flp-17p::eat-4::SL2::GFP* (pMLG017) [27] by digesting it with NheI and KpnI to remove the *eat-4* cDNA sequence. An *egl-4* cDNA fragment containing isoform 2a.1 was PCR-amplified from the plasmid pFG99 *srb-6p::egl-4.2a.1* (a gift from D. Ferkey) [37] using primers containing AvrII and KpnI sites, then ligated downstream of the *flp-17* promoter.

#### AFD

For cell-specific expression in AFD, the plasmid *gcy-8p::egl-4.2a.1::SL2::GFP* (pRFF002) was generated from pRFF001 by digesting pRFF001 with NotI and BamHI to remove the *flp-17* promoter, and then replacing the *flp-17* promoter with the *gcy-8* promoter, which had been cut out of the plasmid AT1_47 *gcy-8p::gcy-8::SL2::mCherry* (a gift from P. Sengupta) [66] using the same enzymes.

#### URX, AQR, and PQR

For cell-specific expression in URX, AQR, and PQR, the plasmid *gcy-36p::egl-4::SL2::GFP* (pRFF003) was generated by first amplifying the *gcy-36* promoter from a *gcy-36p::GCaMP3.0* plasmid [67] with primers containing NotI and BamHI sites, and then ligating it into the *egl-4.2a.1::SL2::GFP* backbone generated from pRFF001 that was previously digested with NotI and BamHI.

#### ASG

For cell-specific expression in ASG, the plasmid *gcy-21p::egl-4.2a.1::SL2::GFP* (pRFF006) was generated by first amplifying the *gcy-21* promoter from plasmid pNTN123 *gcy-21p::YC3.60* (a gift from A. Kuhara) [68] using primers containing NotI and BamHI sites, and then inserting it into the *egl-4.2a.1::SL2::GFP* backbone generated from pRFF001.

#### ASJ

For cell-specific expression in ASJ, the plasmid *ssu-1p::egl-4.2a.1::SL2::GFP* (pAEB001) was generated by PCR-amplifying the *ssu-1* promoter from a plasmid containing the *ssu-1* promoter in the pPD95.77 vector (a gift from A. Miranda-Vizuete) [69] using primers containing NotI and BamHI sites, and then cloning the promoter into the *egl-4.2a.1::SL2::GFP* backbone generated from pRFF001.

#### ASK

For cell-specific expression in ASK, the plasmid *sra-9p::egl-4.2a.1::SL2::GFP* (pRFF007) was generated by excising the *sra-9* promoter from the *sra-9p::tax-4::SL2::GFP* plasmid (a gift from E. Glater) [70, 71] using NotI and BamHI, and then ligating it into the *egl-4.2a.1::SL2::GFP* backbone generated from pRFF001. The pRFF007 plasmid was then injected at a concentration of 100 ng/µL.

BAG-specific expression of the *egl-4* gain-of-function allele *egl-4*(*mg410*) [34] was achieved using the plasmid *flp-17p::egl-4(mg410)::SL2::GFP* (pRFF012). This plasmid was generated by inserting a synthesized *egl-4(mg410)* cDNA containing AvrII and KpnI sites (Twist Bioscience) into pMLG017 that had been previously digested with NheI and KpnI. The pRFF012 construct was injected into wild-type animals.

BAG-specific expression of *flp-17* was achieved with a *gcy-33p::flp-17::SL2::GFP* plasmid (pRFF011) that was generated by inserting the *gcy-33* promoter, amplified from a *gcy-33p::DsRed* plasmid with primers containing NotI and XmaI sites, into a backbone containing the *flp-17 gene*, which was amplified from genomic DNA. The pRFF011 plasmid was injected into *flp-17(ky185)* animals.

To generate the *flp-17p::FlincG3* construct (pRFF013) for quantifying cGMP levels in BAG, the MKV937 *nlp-1p::FlincG3* plasmid (Addgene plasmid #140508, a gift from M. VanHoven) [47] was digested with NotI and XmaI to remove the *nlp-1* promoter and insert the *flp-17* promoter, which had been cut out of a *flp-17p::flp-17::SL2::GFP* plasmid. The *flp-17p::FlincG3* plasmid was injected into wild-type animals with the *unc-122p::DsRed* construct as a co-injection marker. The coinjection marker was used to select animals in an unbiased manner for imaging and quantification.

All primer sequences for PCR-amplification of promoters are listed in Table S2. Amplifications were performed with Platinum Taq High Fidelity (ThermoFisher Scientific) to ensure sequence accuracy. Sequence accuracy was confirmed with whole-plasmid sequencing (Plasmidsaurus).

### CO_2_ chemotaxis assays

Animals were prepared for CO_2_ chemotaxis assays essentially as previously described [26–29], with some modifications.

#### Preparation of well-fed animals raised at low CO_2_

Animals were maintained on OP50 lawns for 3 days before performing experiments. Animals were washed off plates with M9 buffer [62] and transferred to a 65 mm Syracuse watch glass. Animals were allowed to settle before excess M9 was removed. The animals were subsequently washed twice with fresh M9 and once with ddH_2_O. Excess water was removed, and animals were then transferred onto a 10 cm 2% NGM plate [62] for assays using a small piece of Whatman filter paper.

#### Preparation of starved animals raised at low CO_2_

Animals were collected from OP50 plates as described above and placed onto unseeded 2% NGM plates containing an annular ring of Whatman paper dipped in 20 mM copper chloride (CuCl_2_) to prevent the animals from roaming off the plate during starvation [72]. Animals were starved for 3 h. The ring of CuCl_2_ was then removed and the animals were washed off the plate into M9 buffer. The animals were then transferred to an assay plate as described above.

#### Preparation of well-fed animals raised at high CO_2_

Plates were placed in a CO_2_ incubator (Tritech DigiTherm) that was maintained at 23°C and 2.5% CO_2_ for 3 days (*i.e.*, one generation) before experiments were performed, unless otherwise indicated. Plates were kept in the CO_2_ incubator until just prior to processing for the CO_2_ chemotaxis assay. Collections, washes, and transfers were performed as described above for well-fed animals maintained at low CO_2_.

#### CO_2_ chemotaxis assays

Assays were performed essentially as previously described (Fig S1) [26, 28]. Young adult animals were washed off a culturing plate and transferred to an assay plate as described above. Animals were acclimated to the plate for 2 min prior to initiating the chemotaxis assay. A CO_2_ gradient was established by pumping a control gas mixture (21% O_2_, balance N_2_) and a CO_2_ gas mixture (test concentration of CO_2_, 21% O_2_, balance N_2_) into opposite ends of the plate through holes in the lid using a syringe pump (PHD 2000, Harvard Apparatus). Test concentration of CO_2_ was 5% (Fig 2C) or 10% (all other figures). Gas mixtures were delivered through ¼-inch flexible PVC tubing at a flow rate of 2 mL/min (Fig S1A). Assays were run for 20 min, at which point the numbers of animals in each scoring region (Fig S1B) were counted and a chemotaxis index was calculated (Fig S1A). To account for directional bias, the same batch of animals was divided into two groups that were each placed on their own plate, and CO_2_ gradients were established in opposite directions on the two plates. Trials were not counted if the difference between the CI values between the paired plates was 0.9, which was assumed to arise from directional bias.

### Calcium imaging

#### Image acquisition

Calcium imaging was performed using the genetically encoded calcium indicator yellow cameleon YC3.60 [73]. Well-fed young adult animals raised at low CO_2_ were picked individually and moved to 2% NGM plates without food for ∼1 min to remove residual bacteria. Using a size 0 (7 mm) paint brush, animals were transferred to a cover glass with a freshly prepared 2% agarose pad containing 10 mM HEPES. Well-fed animals raised at high CO_2_ were processed as described above, except that the animals were placed in the CO_2_ incubator for 3 days prior to imaging. Separate OP50 plates without animals were also placed in the incubator. These were used for picking single young adult animals 2-3 h before imaging and placed back in the incubator. Each plate was processed separately to minimize time out of the high-CO_2_ environment.

A 15% CO_2_ stimulus was applied using a perfusion chamber with a diameter of 20 mm and a depth of 2.5 mm (Grace Bio-Labs CoverWell, Millipore Sigma) with an adhesive base and two 1.5 mm ports with attached tubing connectors (Grace Bio-Labs, Millipore Sigma) on opposite sides for gas inlet and outlet. The chamber was placed over the agarose pad with the animal and was adhered to the cover glass. Humidified gas was delivered through one port into the chamber through flexible PVC tubing at a flow rate of 30 mL/min using a flow meter (VWR #GR60140AVB-VW) from two gas cylinders fitted with valves controlled by a ValveBank TTL pulse generator. To ensure an airtight seal within the chamber, flexible PVC tubing connected to the gas outlet was immersed in ddH_2_O in a watch glass; water bubbles in the watch glass at the initiation of gas flow confirmed an airtight seal. Total air flow duration was 90 s. An initial air pulse was delivered for 20 s, followed by a 20 s CO_2_ pulse, and then a 50 s air pulse. Calcium imaging was performed using a Zeiss AxioObserver A1 inverted microscope outfitted with a 40x EC Plan-NEOFLUAR objective, a Colibri 7 (Zeiss) for LED fluorescence illumination, a 78 HE ms (1) filter set (BP445/25 + BP510/15, DFT460+520; Zeiss), and a Hamamatsu ORCA-Flash4.0 camera for simultaneous acquisition of CFP and YFP signals. Images were acquired in the YFP and CFP channels at 2 frames/s using Zeiss ZEN 3.4 (Blue edition) software. The emission signals were passed through a Hamamatsu W-View Gemini beam splitter with a CFP/YFP dual filter set.

#### Image processing and data analysis

Calcium traces (normalized and background-subtracted) for each BAG neuron were obtained using Zeiss Zen software (Blue edition) for image analysis and Microsoft Excel for data analysis. Each frame image was analyzed by selecting a region of interest (ROI) that tightly surrounded the BAG soma, and a separate one consisting of a background region off the animal. The same ROI, one for the soma and the other for the background, were propagated to all other frames. The ROI dimensions were the same for all frames but the position of the ROI for the soma was adjusted on each frame, as needed, to correct for small movements in the head of the animal. The average intensity for YFP and CFP of the background region was subtracted from the average intensity for YFP and CFP of the soma. YFP values were adjusted to correct for CFP signal bleed-through, and the YFP/CFP ratio was then calculated. For each of the three conditions under which the animals were raised, all three were tested in parallel across at least 3 days. For quantification of maximum calcium response in each trace, the response period was defined as the time interval beginning with the onset of the CO_2_ pulse and ending 10 s after the offset of the CO_2_ pulse; the % ΔR/R_0_ (max) values were calculated during this response period. Graphs and heatmaps were generated using GraphPad Prism v11. The calcium responses of each animal within the heatmaps were ordered by hierarchical cluster analysis, using Euclidean distance as a similarity measure. Hierarchical cluster analysis was performed using the web-based tool Heatmapper [74, 75].

### Fluorescence microscopy for quantification

#### Preparation of well-fed animals raised at low CO_2_

Well-fed animals were maintained as described above for CO_2_ chemotaxis assays, except that no washing was performed when removing animals from plates. Animals were picked from plates maintained at room temperature (∼23°C).

#### Preparation of starved animals raised at low CO_2_

An annular ring of Whatman paper soaked in 20 mM CuCl_2_ ring was placed onto a 10 cm 2% NGM plate just prior to transferring worms to prevent the starved worms from leaving the plate [72], as described above. After ∼3 h, individual animals were picked for imaging.

#### Preparation of well-fed animals raised at high CO_2_

Animals were placed in a CO_2_ incubator (Tritech DigiTherm) that was maintained at 23°C and 2.5% CO_2_ for 3 days before experiments to obtain 1-day-old adults that experienced high CO_2_ throughout their development. After screening, animals were kept on a 2% NGM plate with OP50 that was stored in the CO_2_ incubator. Five animals were taken out at a time from the plate; the plate was then immediately returned to the incubator to minimize time spent out of the high CO_2_ environment.

#### Image acquisition and analysis

Imaging was performed on a Zeiss AxioObserver microscope equipped with a Colibri 7 for LED fluorescence illumination and a Hamamatsu ORCA-Flash 4.0 camera. For monitoring cGMP levels in BAG neurons (Fig 3), epifluorescence images were acquired with Zen software (Blue edition) using a Plan-APOCHROMAT 20x objective. Images were acquired at 15% light intensity and 40 ms exposure. Animals were prepared as described below for each condition. For all conditions, animals were screened based on expression of a red coelomocyte marker to avoid bias. Single animals (1-day-old adults) were transferred onto an NGM plate without OP50 to remove residual OP50 from the body. Fresh 10% agarose (dissolved in M9 buffer) pads were prepared on the day of imaging. A 2 µL volume of 0.1 µm polystyrene bead suspension (2.5% w/v) (Polysciences, Inc. 0087615) were placed on top of the agarose pads. Animals were transferred using a paintbrush onto the bead suspension droplet and a coverslip was put on top to immobilize them. The coverslip was sealed using nail polish.

Polystyrene beads were used for immobilization during imaging to avoid any interfering effects of anesthetics on cGMP dynamics [47, 76]. For quantification of *flp-17* expression levels in BAG (Fig 5), images were acquired through a Plan-APOCHROMAT 40x objective using Zen software (Blue edition). Animals were paralyzed using 10 mM levamisole on top of 5% Noble agar pads (dissolved in M9 buffer). Images were captured as z-stacks in Zen and average intensity projections were constructed using Fiji [77]. For each image, the mean fluorescence intensity of a background region of interest (ROI) was subtracted from the mean fluorescence intensity of the neuron cell body (Fig 3) or nucleus (Fig 5), and the resulting intensity values were plotted using GraphPad Prism v11. The background ROI was selected to have the same dimensions as the ROI of the cell body and was placed outside of the cell body but inside the worm image.

### Statistical analysis

Statistical tests were performed using GraphPad Prism v11. Statistical tests used are indicated in the figure legends. Normality tests were performed using the D’Agostino and Pearson test; if data were not normally distributed, non-parametric tests were used. Power analyses were performed using G*power [78].

## Supporting information

Frausto et al Supplemental Data

## Data Availability

Raw data is available from GitHub: https://github.com/HallemLab/Frausto_et_al_2026.

## Acknowledgments

We thank Ruhi Patel and Julian Wagner for insightful comments on the manuscript. We thank Mario de Bono, Denise Ferkey, Manabi Fujiwara, Oliver Hobert, Michael Koelle, Ikue Mori, and David Raizen for strains. We thank Denise Ferkey, Elizabeth Glater, Atsushi Kuhara, Antonio Miranda-Vizuete, Matthew Nelson, Piali Sengupta, and Miri VanHoven for plasmids. Some strains were provided by the *Caenorhabditis* Genetics Center (CGC), which is funded by National Institutes of Health Office of Research Infrastructure Programs (P40OD010440).

## References

1. Zjacic N, Scholz M. The role of food odor in invertebrate foraging. Genes Brain Behav. 2022;21(2):e12793. PMID: 34978135

2. Gu H, Zhao F, Liu Z, Cao P. Defense or death? A review of the neural mechanisms underlying sensory modality-triggered innate defensive behaviors. Curr Opin Neurobiol. 2025;92:102977. PMID: 40015135

3. Lenschow C, Mendes ARP, Lima SQ. Hearing, touching, and multisensory integration during mate choice. Front Neural Circuits. 2022;16:943888. PMID: 36247731

4. Hoglen NEG, Manoli DS. Cupid’s quiver: Integrating sensory cues in rodent mating systems. Front Neural Circuits. 2022;16:944895. PMID: 35958042

5. Banerjee N, Hallem EA. The role of carbon dioxide in nematode behavior and physiology. Parasitology. 2020;147:841–54. PMID:

6. Devineni AV, Scaplen KM. Neural circuits underlying behavioral flexibility: insights from *Drosophila*. Front Behav Neurosci. 2021;15:821680. PMID: 35069145

7. Flavell SW, Gordus A. Dynamic functional connectivity in the static connectome of *Caenorhabditis elegans*. Curr Opin Neurobiol. 2022;73:102515. PMID: 35183877

8. Karigo T, Deutsch D. Flexibility of neural circuits regulating mating behaviors in mice and flies. Front Neural Circuits. 2022;16:949781. PMID: 36426135

9. Uddin LQ. Cognitive and behavioural flexibility: neural mechanisms and clinical considerations. Nat Rev Neurosci. 2021;22(3):167–79. PMID: 33536614

10. Siju KP, De Backer JF, Grunwald Kadow IC. Dopamine modulation of sensory processing and adaptive behavior in flies. Cell Tissue Res. 2021;383(1):207–25. PMID: 33515291

11. Hiroki S, Yoshitane H, Mitsui H, Sato H, Umatani C, Kanda S, et al. Molecular encoding and synaptic decoding of context during salt chemotaxis in *C. elegans*. Nat Commun. 2022;13(1):2928. PMID: 35624091

12. Shanahan LK, Kahnt T. On the state-dependent nature of odor perception. Front Neurosci. 2022;16:964742. PMID: 36090268

13. Oram TB, Card GM. Context-dependent control of behavior in *Drosophila*. Curr Opin Neurobiol. 2022;73:102523. PMID: 35286864

14. Vadasz I, Cummins EP, Brotherton DH, Casalino-Matsuda SM, Dada LA, Green O, et al. Sensing molecular carbon dioxide: a translational focus for respiratory disease. Physiol Rev. 2025;105(4):2657–91. PMID: 40668657

15. Bezerra-Santos MA, Benelli G, Germinara GS, Volf P, Otranto D. Smelly interactions: host-borne volatile organic compounds triggering behavioural responses in mosquitoes, sand flies, and ticks. Parasit Vectors. 2024;17(1):227. PMID: 38755646

16. Amundson RG, Davidson EA. Carbon dioxide and nitrogenous gases in the soil atmosphere. J Geochem Explor. 1990;38:13–41. PMID:

17. 17. Oshins C, Michel F, Louis P, Richard TL, Rynk R. The composting process. The Composting Handbook: Academic Press; 2022. p. 51–101.

18. Lan X, Tans P, Thoning KW. Trends in globally-averaged CO_2_ determined from NOAA global monitoring laboratory measurements. Version 2025-09. National Oceanic and Atmospheric Administration (NOAA) Global Monitoring Laboratory, 2025. 10.15138/9N0H-ZH07.

19. Dahake A. The CO_2_ and humidity senses of insects in a changing world. J Exp Biol. 2026;229. PMID: 41668666

20. van Breugel F, Huda A, Dickinson MH. Distinct activity-gated pathways mediate attraction and aversion to CO_2_ in *Drosophila*. Nature. 2018;564(7736):420–4. PMID: 30464346

21. Wasserman S, Salomon A, Frye MA. *Drosophila* tracks carbon dioxide in flight. Curr Biol. 2013;23(4):301–6. PMID: 23352695

22. Zocchi D, Ye ES, Hauser V, O’Connell TF, Hong EJ. Parallel encoding of CO_2_ in attractive and aversive glomeruli by selective lateral signaling between olfactory afferents. Curr Biol. 2022;32(19):4225–39. PMID: 36070776

23. Hallem EA, Sternberg PW. Acute carbon dioxide avoidance in *Caenorhabditis elegans*. Proc Natl Acad Sci USA. 2008;105(23):8038–43. PMID: 18524955

24. Bretscher AJ, Busch KE, de Bono M. A carbon dioxide avoidance behavior is integrated with responses to ambient oxygen and food in *Caenorhabditis elegans*. Proc Natl Acad Sci USA. 2008;105(23):8044–9. PMID: 18524954

25. Sahu AR, Mallick S, Vats A, Bhattacharya A. Distinct molecular mechanisms regulate feeding state-dependent CO_2_ chemotaxis plasticity during different life stages in *Caenorhabditis elegans*. bioRxiv. 2026. PMID: 42051310

26. Rengarajan S, Yankura KA, Guillermin ML, Fung W, Hallem EA. Feeding state sculpts a circuit for sensory valence in *Caenorhabditis elegans*. Proc Natl Acad Sci USA. 2019;116(5):1776–81. PMID: 30651312

27. Guillermin ML, Carrillo MA, Hallem EA. A single set of interneurons drives opposite behaviors in *C. elegans*. Curr Biol. 2017;27(17):2630–9 PMID:

28. Banerjee N, Shih P-Y, Rojas Palato EJ, Sternberg PW, Hallem EA. Differential processing of a chemosensory cue across life stages sharing the same valence state in *Caenorhabditis elegans*. Proc Natl Acad Sci USA. 2023;120(19):e2218023120. PMID:

29. Banerjee N, Rojas Palato EJ, Shih PY, Sternberg PW, Hallem EA. Distinct neurogenetic mechanisms establish the same chemosensory valence state at different life stages in *Caenorhabditis elegans*. G3. 2024;14(2):jkad271. PMID: 38092065

30. Hallem EA, Spencer WC, McWhirter RD, Zeller G, Henz SR, Ratsch G, et al. Receptor-type guanylate cyclase is required for carbon dioxide sensation by *Caenorhabditis elegans*. Proc Natl Acad Sci USA. 2011;108(1):254–9. PMID: 21173231

31. Riedl J, Fieseler C, Zimmer M. Tyraminergic corollary discharge filters reafferent perception in a chemosensory neuron. Curr Biol. 2022;32:1–11. PMID: 35690069

32. L’Etoile ND, Coburn CM, Eastham J, Kistler A, Gallegos G, Bargmann CI. The cyclic GMP-dependent protein kinase EGL-4 regulates olfactory adaptation in *C. elegans*. Neuron. 2002;36(6):1079–89. PMID: 12495623

33. Smith ES, Martinez-Velazquez L, Ringstad N. A chemoreceptor that detects molecular carbon dioxide. J Biol Chem. 2013;288(52):37071–81. PMID: 24240097

34. Hao Y, Xu N, Box AC, Schaefer L, Kannan K, Zhang Y, et al. Nuclear cGMP-dependent kinase regulates gene expression via activity-dependent recruitment of a conserved histone deacetylase complex. PLoS Genet. 2011;7(5):e1002065. PMID: 21573134

35. Hino T, Hirai S, Ishihara T, Fujiwara M. EGL-4/PKG regulates the role of an interneuron in a chemotaxis circuit of *C. elegans* through mediating integration of sensory signals. Genes Cells. 2021;26(6):411–25. PMID: 33817914

36. van der Linden AM, Wiener S, You YJ, Kim K, Avery L, Sengupta P. The EGL-4 PKG acts with KIN-29 salt-inducible kinase and protein kinase A to regulate chemoreceptor gene expression and sensory behaviors in *Caenorhabditis elegans*. Genetics. 2008;180(3):1475–91. PMID: 18832350

37. Krzyzanowski MC, Brueggemann C, Ezak MJ, Wood JF, Michaels KL, Jackson CA, et al. The *C. elegans* cGMP-dependent protein kinase EGL-4 regulates nociceptive behavioral sensitivity. PLoS Genet. 2013;9(7):e1003619. PMID: 23874221

38. Molina-Garcia L, Colinas-Fischer S, Benavides-Laconcha S, Lin L, Clark E, Treloar NJ, et al. Conflict during learning reconfigures the neural representation of positive valence and approach behavior. Curr Biol. 2024;34(23):5470–83 e7. PMID: 39547234

39. Zhang MG, Seyedolmohadesin M, Mercado SH, Tauffenberger A, Park H, Finnen N, et al. Sensory integration of food and population density during the diapause exit decision involves insulin-like signaling in *Caenorhabditis elegans*. Proc Natl Acad Sci USA. 2024;121(40):e2405391121. PMID: 39316052

40. Ghaddar A, Armingol E, Huynh C, Gevirtzman L, Lewis NE, Waterston R, et al. Whole-body gene expression atlas of an adult metazoan. Sci Adv. 2023;9(25):eadg0506. PMID: 37352352

41. Komatsu H, Mori I, Rhee JS, Akaike N, Ohshima Y. Mutations in a cyclic nucleotide-gated channel lead to abnormal thermosensation and chemosensation in *C. elegans*. Neuron. 1996;17(4):707–18. PMID: 8893027

42. Coburn CM, Bargmann CI. A putative cyclic nucleotide-gated channel is required for sensory development and function in *C. elegans*. Neuron. 1996;17(4):695–706. PMID: 8893026

43. Coates JC, de Bono M. Antagonistic pathways in neurons exposed to body fluid regulate social feeding in *Caenorhabditis elegans*. Nature. 2002;419(6910):925–9. PMID: 12410311

44. Bretscher AJ, Kodama-Namba E, Busch KE, Murphy RJ, Soltesz Z, Laurent P, et al. Temperature, oxygen, and salt-sensing neurons in *C. elegans* are carbon dioxide sensors that control avoidance behavior. Neuron. 2011;69(6):1099–113. PMID: 21435556

45. Rojo Romanos T, Petersen JG, Pocock R. Control of neuropeptide expression by parallel activity-dependent pathways in *Caenorhabditis elegans*. Sci Rep. 2017;7:38734. PMID: 28139692

46. Ferkey DM, Sengupta P, L’Etoile ND. Chemosensory signal transduction in *Caenorhabditis elegans*. Genetics. 2021;217(3):iyab004. PMID: 33693646

47. Woldemariam S, Nagpal J, Hill T, Li J, Schneider MW, Shankar R, et al. Using a robust and sensitive GFP-Based cGMP sensor for real-time imaging in intact *Caenorhabditis elegans*. Genetics. 2019;213(1):59–77. PMID: 31331946

48. Galande KK, Cote RH. Roles of cyclic nucleotide phosphodiesterases in signal transduction pathways in the nematode *Caenorhabditis elegans*. Cells. 2025;14(15). PMID: 40801606

49. Aoki I, Shiota M, Tsukada Y, Nakano S, Mori I. cGMP dynamics that underlies thermosensation in temperature-sensing neuron regulates thermotaxis behavior in *C. elegans*. PLoS One. 2022;17(12):e0278343. PMID: 36472979

50. Beets I, Zhang G, Fenk LA, Chen C, Nelson GM, Felix MA, et al. Natural variation in a dendritic scaffold protein remodels experience-dependent plasticity by altering neuropeptide expression. Neuron. 2020;105:106–21. PMID:

51. Krzyzanowski MC, Woldemariam S, Wood JF, Chaubey AH, Brueggemann C, Bowitch A, et al. Aversive behavior in the nematode *C. elegans* is modulated by cGMP and a neuronal gap junction network. PLoS Genet. 2016;12(7):e1006153. PMID: 27459302

52. Lee JI, O’Halloran DM, Eastham-Anderson J, Juang BT, Kaye JA, Scott Hamilton O, et al. Nuclear entry of a cGMP-dependent kinase converts transient into long-lasting olfactory adaptation. Proc Natl Acad Sci USA. 2010;107(13):6016–21. PMID: 20220099

53. Ripoll-Sanchez L, Watteyne J, Sun H, Fernandez R, Taylor SR, Weinreb A, et al. The neuropeptidergic connectome of *C. elegans*. Neuron. 2023;111(22):3570–89 e5. PMID: 37935195

54. Nakano S, Ikeda M, Tsukada Y, Fei X, Suzuki T, Niino Y, et al. Presynaptic MAST kinase controls opposing postsynaptic responses to convey stimulus valence in *Caenorhabditis elegans*. Proc Natl Acad Sci USA. 2020;117(3):1638–47. PMID: 31911469

55. Li Q, Liberles SD. Aversion and attraction through olfaction. Curr Biol. 2015;25(3):R120–9. PMID: 25649823

56. Knaden M, Hansson BS. Mapping odor valence in the brain of flies and mice. Curr Opin Neurobiol. 2014;24(1):34–8. PMID: 24492076

57. Namburi P, Al-Hasani R, Calhoon GG, Bruchas MR, Tye KM. Architectural representation of valence in the limbic system. Neuropsychopharmacology. 2016;41(7):1697–715. PMID: 26647973

58. Tye KM. Neural circuit motifs in valence processing. Neuron. 2018;100(2):436–52. PMID: 30359607

59. Saad MZH, Ryan VW, Edwards CA, Szymanski BN, Marri AR, Jerow LG, et al. Olfactory combinatorial coding supports risk-reward decision making in *C. elegans*. bioRxiv. 2024. PMID: 39484578

60. Zorab JM, Li H, Awasthi R, Schinasi A, Cho Y, O’Loughlin T, et al. Serotonin and neurotensin inputs in the vCA1 dictate opposing social valence. Nature. 2025;642(8066):154–64. PMID: 40307550

61. Kim J, Lee S, Fang YY, Shin A, Park S, Hashikawa K, et al. Rapid, biphasic CRF neuronal responses encode positive and negative valence. Nat Neurosci. 2019;22(4):576–85. PMID: 30833699

62. Stiernagle T. Maintenance of *C. elegans*. 2006 February 11, 2006. In: WormBook [Internet]. WormBook; [1-11]. Available from: www.wormbook.org.

63. Wang L, Graziano B, Bianchi L. Protocols for treating *C. elegans* with pharmacological agents, osmoles, and salts for imaging and behavioral assays. STAR Protoc. 2023;4(2):102241. PMID: 37104092

64. Evans TC. Transformation and microinjection. 2006 April 6, 2006. In: WormBook [Internet]. Pasadena: WormBook; [1-15]. Available from: www.wormbook.org.

65. Miyabayashi T, Palfreyman MT, Sluder AE, Slack F, Sengupta P. Expression and function of members of a divergent nuclear receptor family in *Caenorhabditis elegans*. Dev Biol. 1999;215(2):314–31. PMID: 10545240

66. Takeishi A, Yu YV, Hapiak VM, Bell HW, O’Leary T, Sengupta P. Receptor-type guanylyl cyclases confer thermosensory responses in *C. elegans*. Neuron. 2016;90(2):235–44. PMID: 27041501

67. Carrillo MA, Guillermin ML, Rengarajan S, Okubo R, Hallem EA. O_2_-sensing neurons control CO_2_ response in *C. elegans*. J Neurosci. 2013;33:9675–83. PMID:

68. Takagaki N, Ohta A, Ohnishi K, Kawanabe A, Minakuchi Y, Toyoda A, et al. The mechanoreceptor DEG-1 regulates cold tolerance in *Caenorhabditis elegans*. EMBO Rep. 2020;21(3):e48671. PMID: 32009302

69. Gonzalez-Barrios M, Fierro-Gonzalez JC, Krpelanova E, Mora-Lorca JA, Pedrajas JR, Penate X, et al. Cis- and trans-regulatory mechanisms of gene expression in the ASJ sensory neuron of *Caenorhabditis elegans*. Genetics. 2015;200(1):123–34. PMID: 25769980

70. Macosko EZ, Pokala N, Feinberg EH, Chalasani SH, Butcher RA, Clardy J, et al. A hub-and-spoke circuit drives pheromone attraction and social behaviour in *C. elegans*. Nature. 2009;458(7242):1171–5. PMID: 19349961

71. Worthy SE, Rojas GL, Taylor CJ, Glater EE. Identification of odor blend used by *Caenorhabditis elegans* for pathogen recognition. Chem Senses. 2018;43(3):169–80. PMID: 29373666

72. Campbell JC, Chin-Sang ID, Bendena WG. A *Caenorhabditis elegans* nutritional-status based copper aversion assay. J Vis Exp. 2017;125:e55939. PMID: 28784963

73. Nagai T, Yamada S, Tominaga T, Ichikawa M, Miyawaki A. Expanded dynamic range of fluorescent indicators for Ca^2+^ by circularly permuted yellow fluorescent proteins. Proc Natl Acad Sci USA. 2004;101(29):10554–9. PMID: 15247428

74. Babicki S, Arndt D, Marcu A, Liang YJ, Grant JR, Maciejewski A, et al. Heatmapper: web-enabled heat mapping for all. Nucleic Acids Res. 2016;44(W1):W147–W53. PMID: WOS:000379786800025

75. Kernick K, Woudstra R, Berjanskii M, MacKay S, Wishart DS. Heatmapper2: web-enabled heat mapping made easy. Nucleic Acids Res. 2025;53(W1):W316–W23. PMID: 40322914

76. Cianciulli A, Yoslov L, Buscemi K, Sullivan N, Vance RT, Janton F, et al. Interneurons regulate locomotion quiescence via cyclic adenosine monophosphate signaling during stress-induced sleep in *Caenorhabditis elegans*. Genetics. 2019;213(1):267–79. PMID: 31292211

77. Schindelin J, Arganda-Carreras I, Frise E, Kaynig V, Longair M, Pietzsch T, et al. Fiji: an open-source platform for biological-image analysis. Nat Methods. 2012;9(7):676–82. PMID: 22743772

78. Faul F, Erdfelder E, Lang AG, Buchner A. G*Power 3: a flexible statistical power analysis program for the social, behavioral, and biomedical sciences. Behav Res Methods. 2007;39(2):175–91. PMID: 17695343

