## Supplementary material for "cGMP signaling regulates context-dependent sensory valence in *C. elegans*": Frausto et al Supplemental Data

### Supplemental Figures

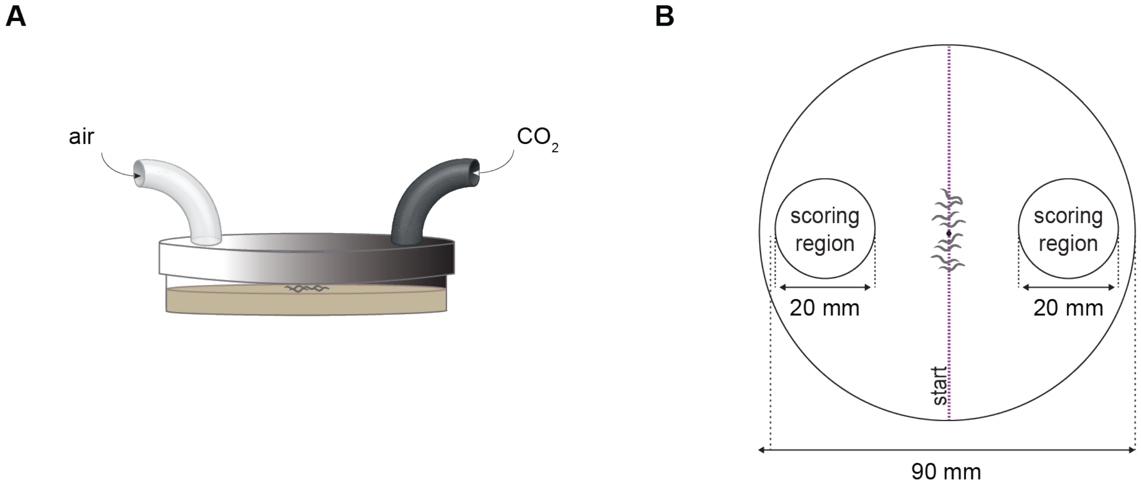


$$\text{chemotaxis index} = \frac{(\text{\# animals at } CO_2 - \text{\# animals at air})}{(\text{\# animals at } CO_2 + \text{air control})}$$

**Fig S1. The $CO_2$ chemotaxis assay.** (**A**) Schematic of the $CO_2$ chemotaxis assay. A $CO_2$ gradient was established across the arena by delivering a gas mixture containing 0% $CO_2$, 21% $O_2$, balance $N_2$ into one side of the plate and a second gas mixture containing the test concentration of $CO_2$, 21% $O_2$, balance $N_2$ into the other side. Young adult hermaphrodites were placed in the center of the arena and allowed to navigate in the $CO_2$ gradient. **(B)** Dimensions of the $CO_2$ assay arena. Animals were placed in the center of the plate (represented by the magenta dashed line) at the start of the assay. After 20 min, the number of animals in each scoring region was counted and the chemotaxis index was calculated using the formula shown. Worms not to scale.

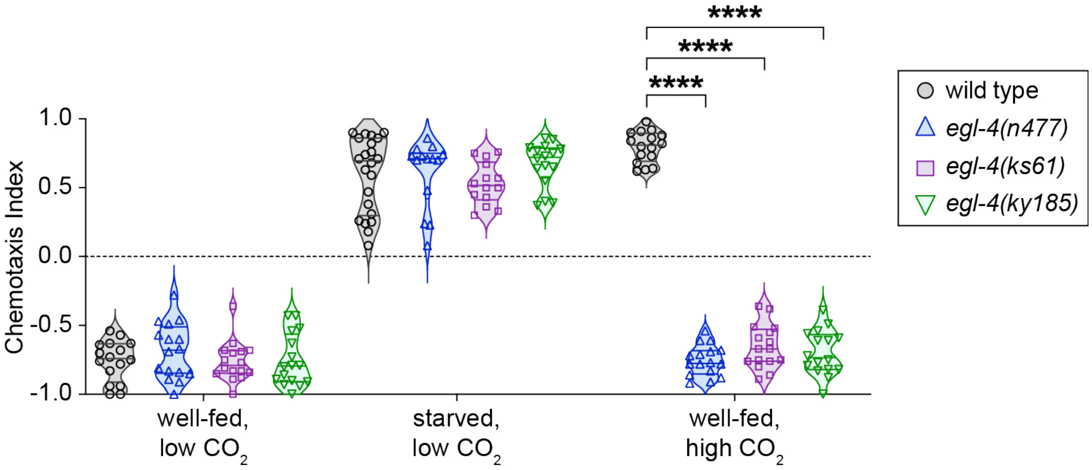


**Fig S2. Confirmation of the *egl-4(lof)* phenotype with multiple independent alleles.** Chemotaxis of wild-type animals and three different *egl-4(lof)* mutants across conditions. ****$p$<0.0001, two-way ANOVA with Dunnett's post-test; only significant comparisons within each condition are shown. n = 14-22 trials per genotype and condition. In the graph, symbols represent individual trials and lines indicate medians and interquartile ranges.

### Supplemental Tables

**Table S1. The list of strains used.** Strains are listed in the order in which they appear. CGC refers to the *Caenorhabditis* Genetics Center.

| Strain | Description | References | Source |
|---|---|---|---|
| N2 Bristol | wild type | Corsi *et al.*, 2015 [1] | CGC |
| MT1074 *egl-4(n479)* | *egl-4* loss-of-function mutant | Trent *et al.*, 1983 [2] | CGC |
| *egl-4(ky185)* | *egl-4* loss-of-function mutant | Hino *et al.*, 2021 [3]; Fujiwara *et al.*, 2022 [4] | M. Fujiwara |
| LX2247 *egl-4(mg410)* | *egl-4* gain-of-function mutant | Hao *et al.*, 2011 [5] | M. Koelle |
| QD87 *egl-4(ky185); Ex[egl-4ap::egl-4.a, myo-3p::GFP]* | *egl-4* rescue in EGL-4-positive cells | Hino *et al.*, 2021 [3] | M. Fujiwara |
| QD86 *egl-4(ky185); Ex[H20p::egl-4.a::GFP(N), myo-3p::GFP]* | *egl-4* pan-neuronal rescue | Hino *et al.*, 2021 [3] | M. Fujiwara |
| QD88 *egl-4(ky185); Ex[tax-4p::egl-4.a, myo-3p::GFP]* | *egl-4* rescue in TAX-4-positive neurons | Hino *et al.*, 2021 [3] | M. Fujiwara |
| EAH447 *egl-4(ky185); bruEx223[gcy-8p::egl-4.2a.1::SL2::GFP]* | *egl-4* rescue in AFD neurons | This study | |
| EAH453 *egl-4(ky185); bruEx227[gcy-21p::egl-4.2a.1::SL2::GFP]* | *egl-4* rescue in ASG neurons | This study | |
| QD92 *egl-4(ky185); Ex[gpa-4p::egl-4.a, myo-3p::GFP]* | *egl-4* rescue in ASI and AWA neurons | Hino *et al.*, 2021 [3] | M. Fujiwara |
| QD90 *egl-4(ky185); Ex[ceh-36p::egl-4.a, myo-3p::GFP]* | *egl-4* rescue in ASE and AWC neurons | Hino *et al.*, 2021 [3] | M. Fujiwara |
| QD96 *egl-4(ky185); Ex[odr-10p::egl-4.a, gcy-10p::egl-4.a; myo-3p::GFP]* | *egl-4* rescue in AWA, AWB, and AWC neurons | Hino *et al.*, 2021 [3] | M. Fujiwara |
| EAH441 *egl-4(ky185); bruEx217[flp-17p::egl-4.2a.1::SL2::GFP]* | *egl-4* rescue in BAG neurons | This study | |
| EAH436 *egl-4(ky185); bruEx212[gcy-36p::egl-4.2a.1::SL2::GFP]* | *egl-4* rescue in URX, AQR, and PQR neurons | This study | |
| EAH458 *egl-4(ky185); bruEx233[ssu-1p::egl-4.2a.1::SL2::GFP]* | *egl-4* rescue in ASJ neurons | This study | |

| | | | |
|---|---|---|---|
| EAH461 *egl-4(ky185); bruEx236[sra-9p::egl-4.2a.1::SL2::GFP]* | *egl-4* rescue in ASK neurons | This study | |
| EAH481 *bruEx245[flp-17p::egl-4(mg410)::SL2::GFP]* | *egl-4(mg410)* overexpression in BAG neurons | This study | |
| EAH498 *bruEx258[flp-17p::FlincG3, unc-122p::DsRed]* | FlincG3 expression in BAG neurons | This study | |
| IK1401 *pde-5(nj49) pde-1(nj57); pde-3(nj59); pde-2(nj58)* | quadruple loss-of-function mutant lacking *pde-1*, *pde-2*, *pde-3*, and *pde-5* | Aoki *et al.,* 2022 [6] | I. Mori |
| IK0616 *pde-1(nj57)* | *pde-1* loss-of-function mutant | Aoki *et al.,* 2022 [6] | I. Mori |
| AX2073 *lin-15(n765ts); dbEx[flp-17p::YC3.60, lin-15(+)]* | yellow cameleon YC3.60 expression in BAG neurons | Kodama-Namba *et al.,* 2013 [7] | M. de Bono |
| EAH417 *egl-4(ky185); dbEx[flp-17p::YC3.60, lin-15(+)]* | yellow cameleon YC3.60 expression in the BAG neurons in the *egl-4(lof)* background | This study | |
| PHX3184 *flp-17(syb3184[flp-17::T2A::3xNLS::GFP])* | strain in which *flp-17* is endogenously tagged | G. Valperga and O. Hobert, in preparation | CGC |
| EAH501 *egl-4(ky185); flp17(syb3184[flp-17::T2A::3xNLS::GFP])* | strain in which *flp-17* is endogenously tagged in the *egl-4(lof)* background | This study | |
| EAH473 *egl-4(ky185); bruEx244[gcy-33p::flp-17::SL2::GFP]* | *flp-17* overexpression in the *egl-4(lof)* background | This study | |
| MT1072 *egl-4(n477)* | *egl-4* loss-of-function mutant | Daniels *et al.,* 2000 [8]; Hirose *et al.,* 2003 [9]; Trent *et al.,* 1983 [2] | CGC |
| FK229 *egl-4(ks61)* | *egl-4* loss-of-function mutant | Hirose *et al.,* 2003 [9] | CGC |

**Table S2. Primer sequences used for molecular biology and genotyping.** Size refers to the length in base pairs of the amplicon produced.

| Oligonucleotide sequence (5' to 3') | Purpose | Sizes (bp) |
|---|---|---|
| F1: GCCTGTTTAGCCACGCCCTA<br>F2: CGACTATTGGGCTCTTGGAAT<br>R: GGTTTCAACGTCCTACTTCTC | genotype *egl-4(ky185)* allele | F1, R (WT): 1230<br>F2, R (WT): 288<br>F1, R (ky185): 453<br>F2, R (ky185): none |
| F: TTATAACCTAGGATGAGCTCTGGGAGCCGTC<br>R: AACATTGGTAC/CCTAGAATCCCTCATCCCATC | amplify *egl-4* cDNA | 2369 |
| F: TTATAACCTAGGATGCTCTCCAAACTAGTGCTCACC<br>R: AACATTGGTACCTTATTTTCCAAAGCGAATGTACTGGCTC | amplify *flp-17* gene | 1377 |
| F: TTATAAGCGGCCGCGATGTTGGTAGATGGGGTT<br>R: ATTCGGGGATCCTGTTGGGTAG | amplify *gcy-36* promoter | 1109 |
| F: TTATAAGC/GGCCGCATCATTCAATGGAATATTTC<br>R: ATTCGGG/GATCCGCATCGTTTCAAC | amplify *trx-1* promoter | 880 |
| F: TTATAAGCGGCCGCCCAGCTCCGCCCCACTAAT<br>R: ATTCGGGGATCCCGGTGGGGTCC | amplify *ssu-1* promoter | 340 |
| F: TTATAAGCGGCCGCTAAACTGGGAGTGAAAGCATCTC<br>R: ATTCGGGGATCCAGCAGAATAATATGAAAATGAAATTTATTTAG | amplify *gcy-21* promoter | 1463 |
| F: TTATAAGCGGCCGCTATCCTGGTCATATCAACTTTCCAGCA<br>R: ATTCGGCCCGGGTTTGCGGGTTGATTTCTGCTAGAAG | amplify *gcy-33* promoter | 1027 |

**Supplemental References**

1. Corsi AK, Wightman B, Chalfie M. A transparent window into biology: a primer on *Caenorhabditis elegans*. Genetics. 2015;200(2):387-407. PMID: 26088431

2. Trent C, Tsuing N, Horvitz HR. Egg-laying defective mutants of the nematode *Caenorhabditis elegans*. Genetics. 1983;104(4):619-47. PMID: 11813735

3. Hino T, Hirai S, Ishihara T, Fujiwara M. EGL-4/PKG regulates the role of an interneuron in a chemotaxis circuit of *C. elegans* through mediating integration of sensory signals. Genes Cells. 2021;26(6):411-25. PMID: 33817914

4. Fujiwara M, Sengupta P, McIntire SL. Regulation of body size and behavioral state of *C. elegans* by sensory perception and the EGL-4 cGMP-dependent protein kinase. Neuron. 2002;36(6):1091-102. PMID: 12495624

5. Hao Y, Xu N, Box AC, Schaefer L, Kannan K, Zhang Y, et al. Nuclear cGMP-dependent kinase regulates gene expression via activity-dependent recruitment of a conserved histone deacetylase complex. PLoS Genet. 2011;7(5):e1002065. PMID: 21573134

6. Aoki I, Shiota M, Tsukada Y, Nakano S, Mori I. cGMP dynamics that underlies thermosensation in temperature-sensing neuron regulates thermotaxis behavior in *C. elegans*. PLoS One. 2022;17(12):e0278343. PMID: 36472979

7. Kodama-Namba E, Fenk LA, Bretscher AJ, Gross E, Busch KE, de Bono M. Cross-modulation of homeostatic responses to temperature, oxygen and carbon dioxide in *C. elegans*. PLoS Genet. 2013;9(12):e1004011. PMID: 24385919

8. Daniels SA, Ailion M, Thomas JH, Sengupta P. *egl-4* acts through a transforming growth factor-beta/SMAD pathway in *Caenorhabditis elegans* to regulate multiple neuronal circuits in response to sensory cues. Genetics. 2000;156(1):123-41. PMID: 10978280

9. Hirose T, Nakano Y, Nagamatsu Y, Misumi T, Ohta H, Ohshima Y. Cyclic GMP-dependent protein kinase EGL-4 controls body size and lifespan in *C. elegans*. Development. 2003;130(6):1089-99. PMID: 12571101
